# DNA methylation and histone H1 cooperatively repress transposable elements and aberrant intragenic transcripts

**DOI:** 10.1101/527523

**Authors:** Jaemyung Choi, David B. Lyons, M. Yvonne Kim, Jonathan D. Moore, Daniel Zilberman

## Abstract

DNA methylation and histone H1 mediate transcriptional silencing of genes and transposable elements, but how they interact is unclear. In plants and animals with mosaic genomic methylation, functionally mysterious methylation is also common within constitutively active housekeeping genes. Here we show that H1 is enriched in methylated sequences, including genes, of *Arabidopsis thaliana*, yet this enrichment is independent of DNA methylation. Loss of H1 disperses heterochromatin, globally alters nucleosome organization, and activates H1-bound genes, but only weakly de-represses transposable elements. However, H1 loss strongly activates transposable elements hypomethylated through mutation of DNA methyltransferase *MET1*. Loss of H1 also activates antisense transcripts within demethylated genes. Our results demonstrate that H1 and DNA methylation cooperatively maintain transcriptional homeostasis by silencing transposable elements and aberrant intragenic transcripts. Such functionality plausibly explains why DNA methylation, a well-known mutagen, has been maintained within coding sequences of crucial plant and animal genes.

**Highlights:**

- Histone H1 is enriched in methylated DNA independently of methylation
- Loss of H1 activates genes, alters nucleosome organization and disperses heterochromatin
- DNA methylation and H1 jointly silence transposons
- DNA methylation and H1 cooperatively suppress intragenic antisense transcripts

## Introduction

Cytosine DNA methylation is an epigenetic modification that protects genome integrity by repressing transposable elements (TEs) in plants, vertebrates, fungi, and likely other eukaryotic groups (Du et al., 2015; Jones, 2012; Zemach and Zilberman, 2010). Methylation of regulatory sequences, such as promoters and enhancers, also causes gene silencing in plants and vertebrates (Hon et al., 2013; Jones, 2012; Schübeler, 2015; Zhang et al., 2018a). Proper patterns of DNA methylation are essential for plant and animal development (Iurlaro et al., 2017; Kawashima and Berger, 2014; Schmidt et al., 2015; Smith and Meissner, 2013), and their disruption is linked with cancer and other serious diseases (Jones et al., 2016; Rasmussen and Helin, 2016; Robertson, 2005).

Despite its association with transcriptional repression, DNA methylation is common within active genes (Bewick and Schmitz, 2017; Feng et al., 2010; Jones, 2012; Zemach et al., 2010). Vertebrate intragenic methylation occurs in the context of a globally methylated genome, in which methylation is the default state and some sequences, including active promoters and enhancers, are protected from methylation (Schübeler, 2015; Suzuki and Bird, 2008). Mammalian genes are co-transcriptionally hypermethylated by the *de novo* methyltransferase Dnmt3 (Baubec et al., 2015; Morselli et al., 2015; Neri et al., 2017), and – likely due to this activity – mammalian intragenic methylation is positively correlated with gene expression (Lister et al., 2009; Yang et al., 2014). Several functions have been ascribed to mammalian intragenic methylation, including the silencing of intragenic promoters and enhancers (Deaton et al., 2011; Hellman and Chess, 2007; Kulis et al., 2012; Maunakea et al., 2010; Yang et al., 2014), regulation of splicing (Lev Maor et al., 2015; Maunakea et al., 2013; Shukla et al., 2011), and suppression of aberrant transcripts (Neri et al., 2017). However, the latter function has been questioned, because strong DNA methylation loss in mouse embryonic stem cells was not associated with activation of intragenic transcription (Teissandier and Bourc’his, 2017).

In contrast to vertebrates, plant and invertebrate genomes usually have mosaic methylation patterns, with specific sequences targeted for methylation (Schübeler, 2015; Suzuki and Bird, 2008). In flowering plants, TEs and gene bodies are preferentially methylated (Bewick and Schmitz, 2017; Lister et al., 2008; Takuno and Gaut, 2013; Zemach and Zilberman, 2010). TEs are affected by multiple pathways that mediate methylation in three sequence contexts (CG, CHG and CHH, where H ≠ G), whereas only CG sites are methylated in genes (Feng et al., 2010; Zemach et al., 2010; Zhang et al., 2018a). Invertebrates preferentially or exclusively methylate gene bodies, and essentially all methylation is in the CG context (Bewick et al., 2017; Dixon et al., 2016; Feng et al., 2010; Suzuki et al., 2007; Zemach et al., 2010). Plant and invertebrate intragenic methylation is not directly linked to transcription, but is instead a largely invariant genomic feature that decorates the same sets of genes across developmental stages and tissues (Bartels et al., 2018; Libbrecht et al., 2016; Suzuki et al., 2013; Zemach et al., 2018). This phenomenon, commonly called gene body methylation (gbM), is concentrated in the exons of evolutionarily conserved, stably and constitutively expressed genes, and is maintained by the methyltransferase Dnmt1 (called MET1 in plants) (Bewick and Schmitz, 2017; Bewick et al., 2018; Cokus et al., 2008; Dixon et al., 2016; Lister et al., 2008; Sarda et al., 2012; Takuno and Gaut, 2012, 2013; Zemach and Zilberman, 2010). GbM has been proposed to regulate splicing and transcriptional elongation, and to suppress intragenic transcriptional initiation (Hunt et al., 2013; Regulski et al., 2013; Suzuki et al., 2007; To et al., 2015; Tran et al., 2005; Zilberman et al., 2007). However, loss of gbM in the flowering plant *Arabidopsis thaliana* and the insect *Oncopeltus fasciatus* is not associated with transcriptional changes (Bewick et al., 2016, 2018; Kawakatsu et al., 2016), and gbM is commonly thought to be a nonfunctional byproduct of TE methylation (Bewick and Schmitz, 2017; Bewick et al., 2016; Teixeira and Colot, 2009). Overall, the functions of gbM – if any exist – remain mysterious (Zhang et al., 2018a; Zilberman, 2017).

A potential explanation for the mystery and controversy surrounding gbM – and intragenic methylation in general – is that its function is subtle and shared with other chromatin factors. This hypothesis was tested in the flowering plant *Arabidopsis thaliana* by simultaneously eliminating gbM and trimethylation of lysine 36 of histone H3, a histone modification that suppresses cryptic intragenic transcripts and regulates splicing (Carrozza et al., 2005; Wagner and Carpenter, 2012), but no transcriptional or RNA processing defects attributable to gbM were found (Bewick et al., 2016). A plausible alternate candidate is histone H1, a protein that binds nucleosomes and the intervening linker DNA (Fyodorov et al., 2018; Hergeth and Schneider, 2015; Over and Michaels, 2014). Like methylation, H1 is associated with transcriptional silencing but is nevertheless abundant in active plant and animal genes (Krishnakumar et al., 2008; Rutowicz et al., 2015; Torres et al., 2016). There is also evidence that H1 preferentially inhibits transcriptional initiation from methylated DNA *in vitro* (Johnson et al., 1995; Levine et al., 1993). These data suggest that H1 and DNA methylation may act cooperatively, including within gene bodies. However, whether H1 affinity is regulated by DNA methylation is uncertain, with some studies finding preferential association with methylated DNA (Levine et al., 1993; McArthur and Thomas, 1996), and others reporting no preference (Campoy et al., 1995; Hashimshony et al., 2003; Nightingale and Wolffe, 1995). More generally, if and how H1 interacts with methylation *in vivo* to regulate transcription is unknown.

## Results

### Histone H1 is enriched in methylated DNA independently of methylation

To investigate the functional relationship between DNA methylation and H1, we analyzed the genomic distribution of the two major *Arabidopsis* H1 proteins, H1.1 and H1.2 (Over and Michaels, 2014). Consistent with earlier work (Rutowicz et al., 2015), we did not find any salient differences between H1.1 and H1.2 (Figure S1A), and therefore will refer simply to H1. As expected, H1 preferentially associates with linker DNA (Figures 1A and S1B), is concentrated in heavily methylated heterochromatic TEs (Figures 1B, 1C, S1C and Table S1), and is abundant within genes, particularly those with low/no expression (Figures 1D and S1D). H1 is also enriched in methylated genes compared to similarly expressed unmethylated genes (Figures 1E and S1E). Thus, H1 is preferentially associated with methylated DNA in *Arabidopsis*.

**Figure 1.**
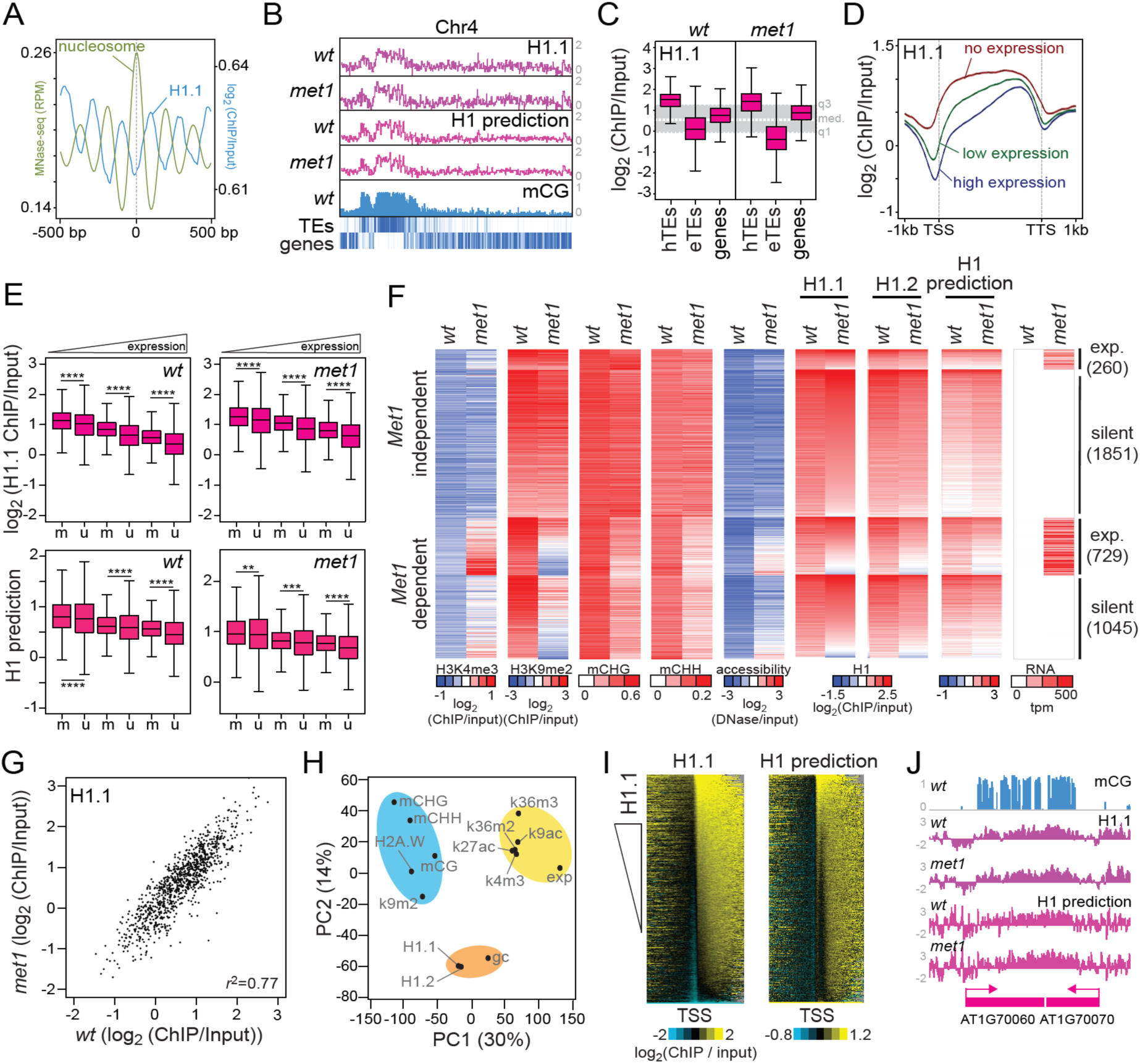
Histone H1 is enriched in methylated DNA independently of methylation. **(A)** Average H1.1 distribution around well-positioned nucleosomes. **(B)** Average H1.1, predicted H1, and CG methylation (mCG) levels are plotted in 50 kb windows along *Arabidopsis* chromosome 4. **(C)** Box plots of H1.1 levels in heterochromatic TEs (hTEs), euchromatic TEs (eTEs), and genes. The shaded area marks the middle 50% of genomic H1.1 levels. **(D)** H1.1 distribution around genes with different expression levels. **(E)** Box plots of H1.1 abundance and predicted H1 abundance in methylated (m) and unmethylated (u) genes in *wt* and *met1* plants. ** is *p* < 0.01, *** is *p* < 0.001, **** is *p* < 0.0001, Student’s *t*-test. **(F)** Heat maps of indicated chromatin features and expression in MET1-independent and dependent hTEs. **(G)** Average H1.1 levels of 1 kb windows in *wt* and *met1*. 1000 windows were randomly selected for plotting. “*r*” is Pearson’s correlation coefficient. Principal component analysis of H1.1, H1.2, histone H3 lysine modifications, histone variant H2A.W, DNA methylation (mCG, mCHG, mCHH), GC content (gc) and gene expression (exp). Linear model prediction of H1 distribution around the transcriptional start sites (TSS) of genes in comparison to actual H1.1 distribution. **(J)** Example of actual H1.1 and predicted H1 distribution at gene-body methylated genes in *wt* and *met1*. **(C and E)** Whiskers indicate 1.5X interquartile range (IQR).

To determine if DNA methylation affects H1 binding, we analyzed H1 distribution in *met1* mutant plants. The *met1* mutation eliminates CG methylation, which includes the entirety of gbM (Cokus et al., 2008; Lister et al., 2008). Many TEs also lose non-CG methylation and dimethylation of lysine 9 of histone H3 (H3K9me2), a mark of heterochromatin (Feng and Michaels, 2015), and become more accessible to DNase I (MET1-dependent TEs, Figure 1F) (Deleris et al., 2012; Zhang et al., 2018b). We find that H1 distribution is well-correlated between wild type (*wt*) and *met1* (Figures 1B, 1C, 1F, 1G and S1C, S1F). Genes methylated in *wt* retain higher levels of H1 in *met1* plants (Figures 1E and S1E), as do TEs (Figures 1C and S1C). Therefore, H1 binding is largely independent of DNA methylation. However, some TEs do lose H1 in *met1* mutants, particularly MET1-dependent TEs that are transcriptionally activated (Figures 1F and S1G).

To determine how H1 distribution is regulated, we analyzed H1 association with various genomic features using principal component analysis. H1 associates most strongly with GC content (Figure 1H) independently of nucleosome density (Figure S1H). Methylated genes have elevated GC content (Figure S1I), which likely explains their H1 enrichment (Figures 1E and S1E). H1 also associates with heterochromatic features, including DNA methylation and H3K9me2, along the first principal component (Figure 1H), and is well-separated from features of active transcription (such as H3K4me3, Figure 1H). Based on these results, we used GC content and *wt* H3K9me2 and H3K4me3 data to create a linear regression model of H1 abundance. This model accurately describes *wt* H1 distribution across genes and TEs (Figures 1B, 1F, 1I, 1J, and S1J-L), including H1 enrichment in methylated genes (Figures 1E and S1 K). More importantly, our model successfully predicts *met1* H1 distribution (Figures 1B, 1E, 1F, 1J and S1J-L), including the loss of H1 from a subset of MET1-dependent (*i.e.* heterochromatin-deficient) TEs (Figures 1F and S1G). Thus, H1 enrichment can largely be explained by sequence composition, transcription, and heterochromatin, whereas DNA methylation has little, if any direct effect on H1 binding.

### Histone H1 mediates global nucleosome organization

The distribution of *Arabidopsis* H1 (heterochromatin > silent genes > active genes, Figures 1B-D) corresponds to an unexplained phenomenon in the human genome, in which genes expressed at higher levels have shorter average nucleosome repeat length (NRL) than less expressed genes, which have shorter average NRLs than heterochromatin (Valouev et al., 2011). Because loss of H1 is known to reduce the average NRL in animal genomes (Baldi et al., 2018; Fan et al., 2003; Woodcock et al., 2006), we tested the hypothesis that H1 causes the observed NRL differences by comparing nucleosome spacing between plants with defective *H1.1* and *H1.2* genes (*h1* mutants) (Zemach et al., 2013) and *wt* controls.

As in the human genome, *Arabidopsis* genes expressed at higher levels have shorter average NRL in *wt* (Figure 2A). The NRL of less-expressed, H1-rich genes decreases substantially in *h1* mutants (Figures 2A-C and S2A). In the most H1-rich genes, mean NRL is reduced from 185 bp to 169 bp (Figures 2B and 2C), which is consistent with the ability of H1 to protect about 20 bp of DNA *in vitro* (Simpson, 1978). Consistent with published results (Zhang et al., 2015), we find that heterochromatic TEs have longer average NRLs than more euchromatic TEs (Figure 2D), with the longest NRLs in the most H1-rich TEs (Figure 2E). Loss of H1 reduces mean NRL of H1-rich TEs by 22 bp (from 189 bp to 167 bp), causing TEs in *h1* mutants to have similar NRLs regardless of their *wt* H1 levels (Figures 2E, 2F and S2B). The average NRLs of TEs and silent genes are also very similar in *h1* plants (Figure S2C). Thus, H1 is an important global regulator of nucleosome placement that accounts for much of the average NRL difference across the *Arabidopsis* genome.

**Figure 2.**
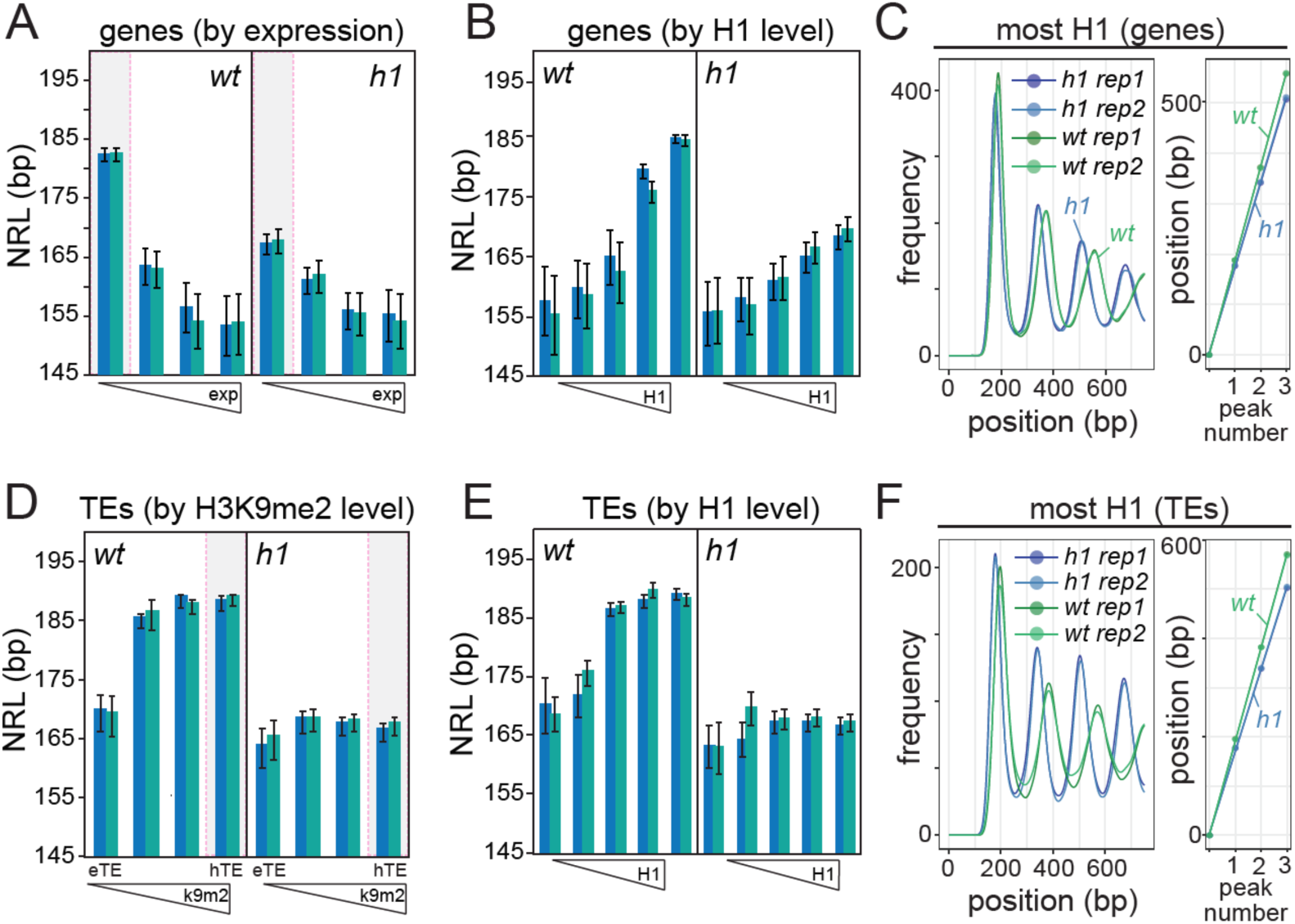
Loss of H1 alters nucleosome organization. **(A)** Nucleosome repeat length (NRL) calculations for two biological replicates at genes grouped by their expression in *wt*. Phasograms and linear regressions for the positions of nucleosome peaks at the genes with lowest expression (shaded area) are in Figure S2A. **(B and E)** NRL calculations for two biological replicates at genes and TEs (E) grouped by H1 enrichment. **(C and F)** Phasograms and linear regressions for the positions of nucleosome peaks at H1-rich genes (C) and TEs (F). **(D)** NRL calculations for two biological replicates at TEs grouped by their H3K9me2 level in *wt*. Phasograms and linear regressions for the positions of nucleosome peaks at H3K9me2-rich TEs (shaded area) are in Figure S2B. **(A-B and D-E)** Error bars indicate standard error.

### Loss of H1 activates transcription and disperses heterochromatin

Histone H1 mediates gene silencing (Fyodorov et al., 2018; Krishnakumar et al., 2008; Torres et al., 2016) and the 3D configuration of heterochromatin (Cao et al., 2013; Lu et al., 2009) in animals, but the effects of H1 on transcription and higher order chromatin organization in plants are unknown. To understand how H1 regulates transcription, we determined mRNA levels using RNA-seq in *h1* mutants in comparison to *wt*. Most of the significantly mis-regulated genes in leaves and seedlings (707/990, 71%) are upregulated in *h1* plants (Figures 3A, 3B and Table S2). These genes are H1-enriched (Figures 3C, 3D and S3A) and are transcribed at low levels in *wt* (Figure 3E), indicating that H1 can repress gene transcription in plants. However, given that H1 is generally abundant in *Arabidopsis* genes (Figures 1C and 1D), its effects on transcription are fairly selective, as is the case in animals (Fan et al., 2005). Several gene ontology functional categories are overrepresented among the upregulated genes, most notably categories related to the meiotic cell cycle and meiotic recombination (Figure S3B and Table S2). Upregulated genes include ASYNAPTIC (ASY) 1 and 4, and other key meiotic genes (Figures 3B, 3F and Table S2) (Chambon et al., 2018; Mercier et al., 2015). H1 is depleted in male and female *Arabidopsis* meiocytes (She and Baroux, 2015; She et al., 2013), suggesting H1 removal may help to activate the meiotic transcriptional program. Genes involved in epigenetic silencing are also preferentially upregulated (Figure S3B and Table S2). Curiously, one of these is ARGONAUTE9 (Figure 3B), which is involved in meiocyte specification (Olmedo-Monfil et al., 2010), further reinforcing the link between H1 and meiocyte fate and function.

**Figure 3.**
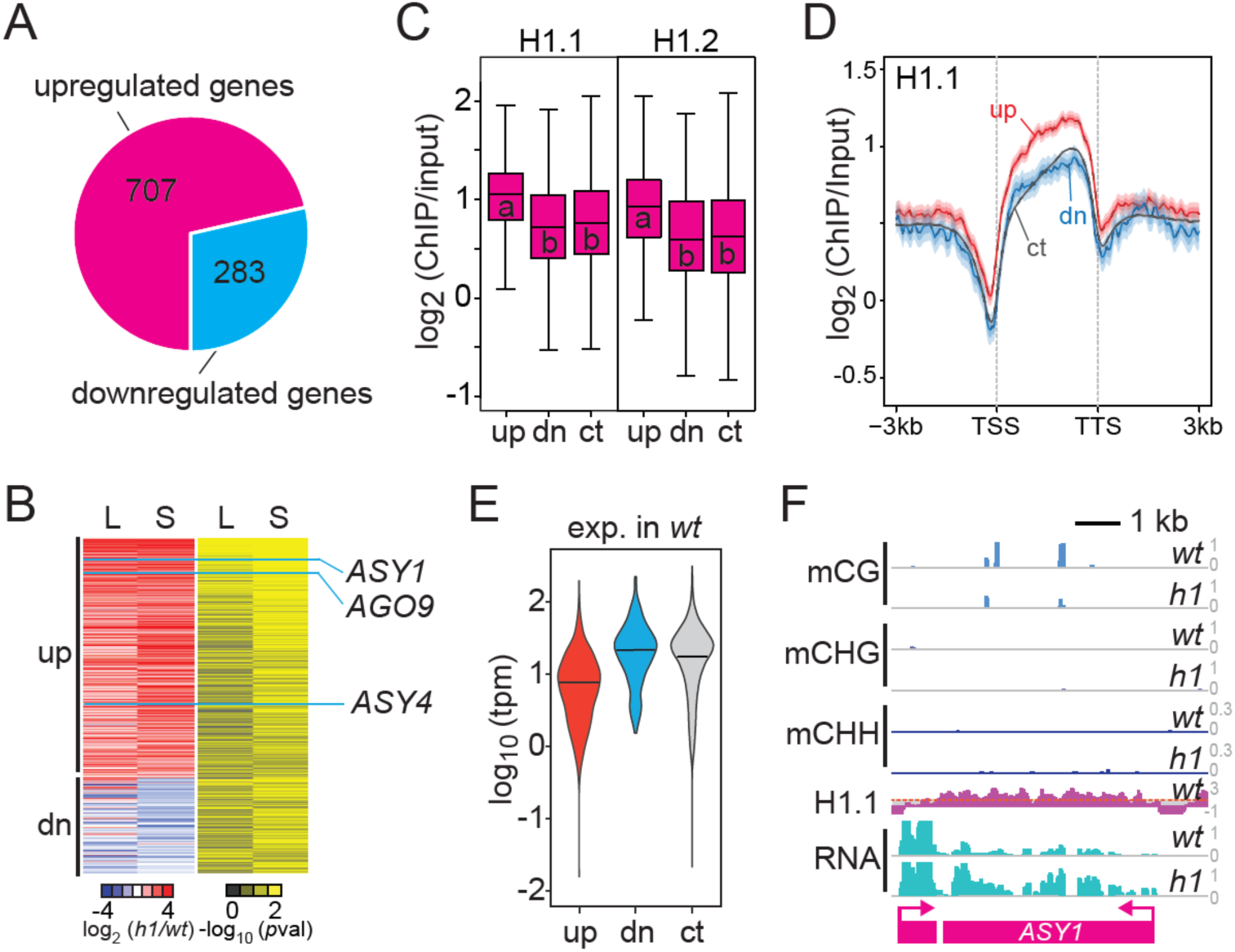
Loss of H1 activates H1-rich meiosis-related genes. **(A)** Number of up- and down- regulated genes in *h1* plants vs. *wt*. **(B)** Heat maps of respective gene expression fold change vs. *wt* in *h1* plants in leaves (L) and seedlings (S). Adjusted *p-*values for the expression change are shown. **(C)** Box plots of *wt* H1 levels in genes upregulated (up), down-regulated (dn) and unchanged (control; ct) in *h1* plants. “a” and “b” are significantly different (*p* < 0.01; ANOVA). Whiskers indicate 1.5X IQR. **(D)** Distribution of H1 around control, up- and down-regulated genes. Shaded areas represent 95% confidence intervals. **(E)** Violin plots of gene expression in *wt* plants for control (ct), up- and down (dn)-regulated genes in (A). **(F)** An example meiosis-related gene transcriptionally activated in *h1* plants. The dashed red line indicates average H1 level at genes.

In *wt Arabidopsis*, heterochromatin is arranged into large DAPI-stained foci called chromocenters (Figures 4A and S4A) (Ascenzi and Gantt, 1999; Fransz et al., 2002). We find that in *h1* mutants nuclei typically have few chromocenters and many smaller DAPI-stained foci (Figures 4A, 4B and S4A), indicating that the role of H1 in heterochromatin organization is generally conserved. However, despite the dispersal of chromocenters, only 32 TEs are significantly upregulated in *h1* plants (Table S2). An important consideration is that H1 is a major regulator of *Arabidopsis* DNA methylation: depletion of H1 increases methylation of some TEs in all sequence contexts, and reduces methylation of other TEs (Lyons and Zilberman, 2017; Rutowicz et al., 2015; Wierzbicki and Jerzmanowski, 2005; Zemach et al., 2013). Most of the upregulated TEs (29, 91%) are hypomethylated in *h1* mutants (Figures 4C-E and S4B), suggesting that some of the observed activation is not directly caused by H1 depletion. Furthermore, heterochromatin does not become substantially more accessible to DNase I in *h1* mutants (Figure 4F). Thus, although heterochromatin is altered (Figures 2D-F) and dispersed (Figures 4A, 4B and S4A) by lack of H1, heterochromatic functionality remains largely intact.

**Figure 4.**
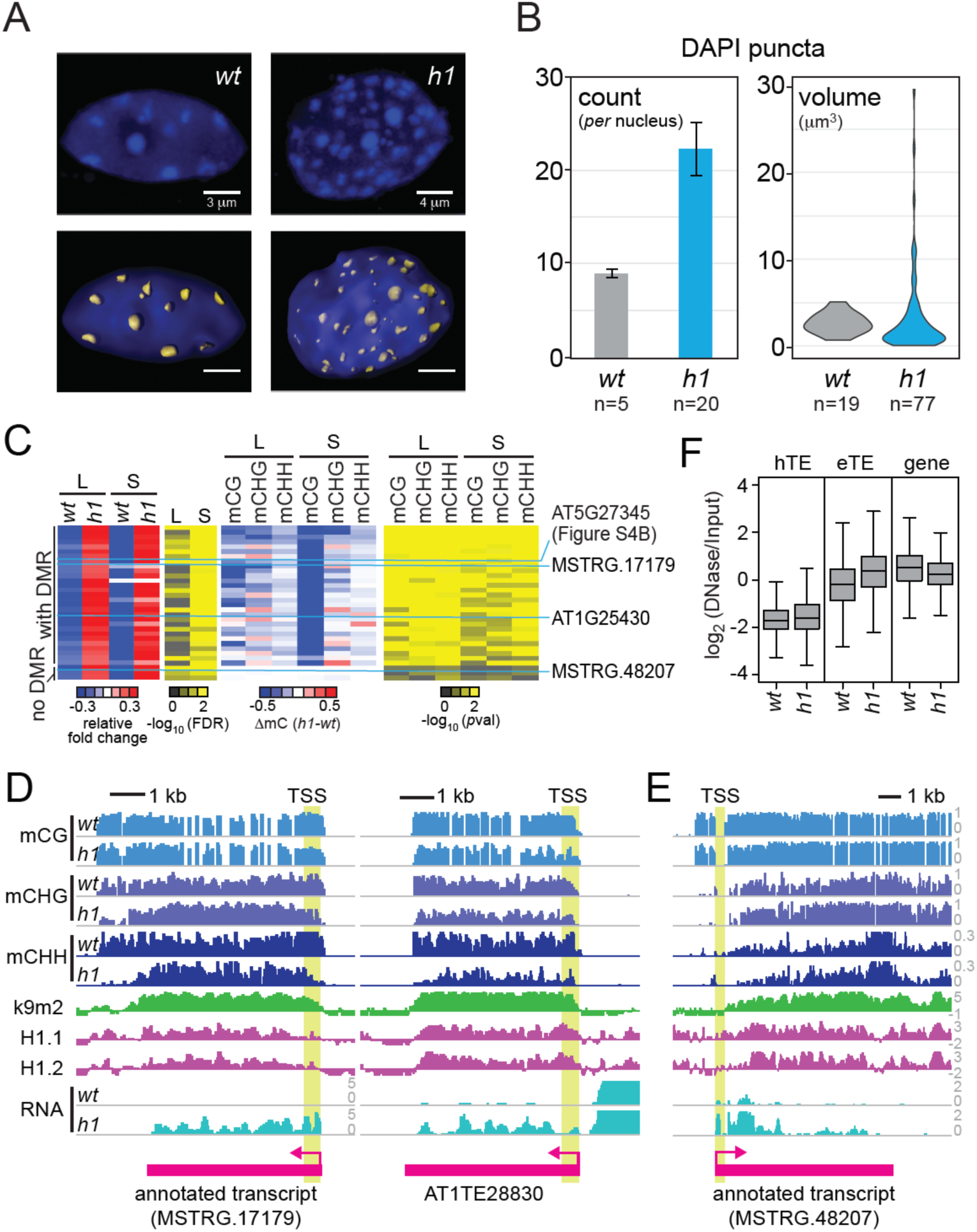
Loss of H1 disperses heterochromatin and weakly activates TEs. **(A)** Nuclei of *wt* and *h1* plants stained with DAPI. DAPI puncta volumes are highlighted in yellow. **(B)** Number per nucleus and volume of DAPI puncta in *wt* and *h1* plants. Note that blurred chromocenter boundaries in *h1* mutants cause some to be assigned very large volumes. The number of nuclei (left) and DAPI puncta (right) used to generate the data were indicated. **(C)** Heat maps of expression in leaves (L) and seedlings (S) with corresponding FDR (*h1* vs. *wt*) and TSS-proximal DNA methylation change of TEs in *h1* plants vs. *wt*. *p*-value for methylation change was calculated with Fisher’s exact test. **(D)** Examples of upregulated TEs in *h1* plants. Note DNA methylation loss near TSS (highlighted in yellow). **(E)** An example of upregulated TE in *h1* plants without DNA methylation loss near TSS (highlighted in yellow). **(F)** Box plots of DNA accessibility of hTEs, eTEs, and genes in *wt* and *h1*. Whiskers indicate 1.5X IQR.

### DNA methylation and H1 cooperatively silence TEs

Limited TE activation in *h1* plants contrasts with the extensive TE upregulation in H1-deficient *Drosophila melanogaster* (Iwasaki et al., 2016; Lu et al., 2013), but is similar to the effects of H1 depletion on mouse cells, which show few changes in TE expression (Fan et al., 2005). TEs are silenced by DNA methylation in mouse and *Arabidopsis* (Law and Jacobsen, 2010), whereas *Drosophila* lacks cytosine methylation (Raddatz et al., 2013; Zemach et al., 2010), suggesting that the effects of H1 loss may be masked by methylation. To test this hypothesis, we investigated TE expression in *h1met1* compound mutants and *met1* controls.

Because H1 can locally raise or lower DNA methylation (Rutowicz et al., 2015; Zemach et al., 2013), we first identified active TEs with unchanged CHG and CHH methylation near the transcriptional start site (TSS) between *met1* and *h1met1*, which turned out to be mostly TEs that are fully demethylated in both genotypes (Figures 5A, 5B, S5A and Table S3). Unsupervised clustering separated these TEs into two groups, both of which become much more accessible to DNase I in *met1* and remain so in *h1met1* (Figures 5A and S5B). The larger group (nc; 272 TEs) is expressed similarly in *met1* and *h1met1* (Figure 5A). However, the smaller group of TEs (up1; 134) is strongly upregulated in *h1met1* (Figures 5A and S5C). These TEs are significantly H1- enriched in *met1* compared to TEs with unchanged expression, especially around the TSS (Figures 5C, 5D and S5D). These TEs are thus apparently transcriptionally repressed by H1 and DNA methylation. Loss of methylation alone leads to mild activation and locus accessibility, but loss of both methylation and H1 is required to create a highly active state (Figures 5A and S5B).

**Figure 5.**
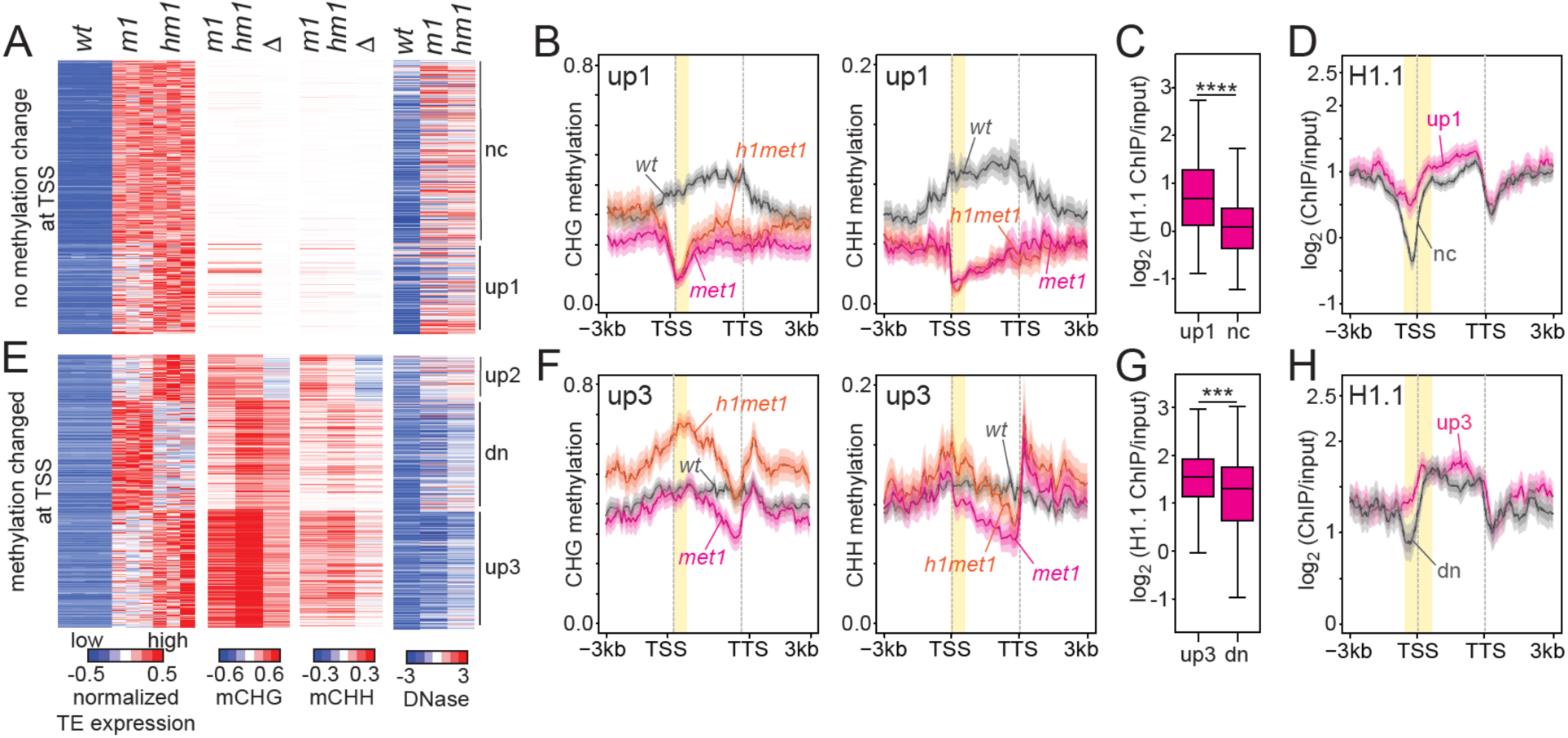
DNA methylation and H1 cooperatively silence TE expression. **(A and E)** Expression, TSS-proximal DNA methylation and DNA accessibility of hTEs in *wt*, *met1* and *h1met1* plants. TEs with unchanged TSS non-CG methylation in *h1met1* compared to *met1* are shown in (A) and TEs with altered non-CG methylation at the TSS are shown in (E). **(B and F)** Average CHG and CHH methylation around TEs in the up1 (B) and up3 (F) clusters. Methylation from the TSS- proximal regions shaded in yellow is shown in (A) and (E). **(C-D)** Box plots of *met1* H1 abundance and H1 distribution (D) in TEs in the nc and up1 clusters from (A). H1 from the TSS-proximal region shaded in yellow in (D) is shown in (C). **** is *p* < 0.0001, Student’s *t*-test. **(G-H)** Box plots of *met1* H1 abundance (G) and H1 distribution (H) in TEs in the dn and up3 clusters from (E). H1 from the TSS-proximal region shaded in yellow in (H) is shown in (G). *** is *p* < 0.001, Student’s *t*-test. **(C and G)** Whiskers indicate 1.5X IQR.

Analysis of TEs with altered TSS-proximal CHG or CHH methylation produced three groups, all of which maintain robust non-CG methylation in *met1* and *h1met1* (Figure 5E and Table S3). All three groups are more accessible to DNase I in *h1met1* compared to *met1* (Figures 5E and S5B), indicating that loss of H1 increases the accessibility of partially demethylated sequences. The first group (up2) is comprised of 63 TEs upregulated and hypomethylated in *h1met1* compared to *met1* (Figures 5E and S5E). The second group (dn) consists of 147 TEs downregulated and hypermethylated in *h1met1* (Figures 5E and S5F). Thus, both groups are likely differentially expressed between *met1* and *h1met1* due to changes in DNA methylation. However, the third – and largest – group (up3; 159 TEs) is upregulated in *h1met1* despite substantial hypermethylation (Figures 5E, 5F and S5G). Compared to the dn group, these TEs are significantly H1-enriched (Figures 5G, 5H and S5D). TEs in the up3 group, like those in group up1 described above, are mildly activated in *met1* and require loss of H1 for strong expression (Figure 5E). These data indicate that DNA methylation and H1 jointly create an inaccessible chromatin state that silences TEs.

### DNA methylation and H1 cooperatively suppress intragenic antisense transcripts

The ability of H1 and DNA methylation to cooperatively repress transcription, and enrichment of H1 within methylated genes (Figure 1E), are consistent with the possibility that methylation and H1 act together to reduce aberrant intragenic transcripts, a long-hypothesized function of gbM (Tran et al., 2005; Zilberman et al., 2007). To test this hypothesis, we identified antisense intragenic transcripts with significantly altered expression in *h1, met1* and *h1met1* mutants in comparison to *wt*. We identified 17 such transcripts in *h1*, most of which (15, 88%) are upregulated (Figure 6A). This result is consistent with the hypothesis that H1 suppresses intragenic transcription, but the small number of identified transcripts limits confidence and prevents further analysis. We found many more mis-expressed antisense transcripts in *met1* (468), and yet more in *h1met1* (1152; Figure 6A). In *met1*, only 32% of transcripts (149) are upregulated (Figure 6A). In contrast, 91% of transcripts (1044) are upregulated in *h1met1* (Figure 6A), supporting the hypothesis that intragenic transcription is de-repressed when DNA methylation and H1 are simultaneously absent.

**Figure 6.**
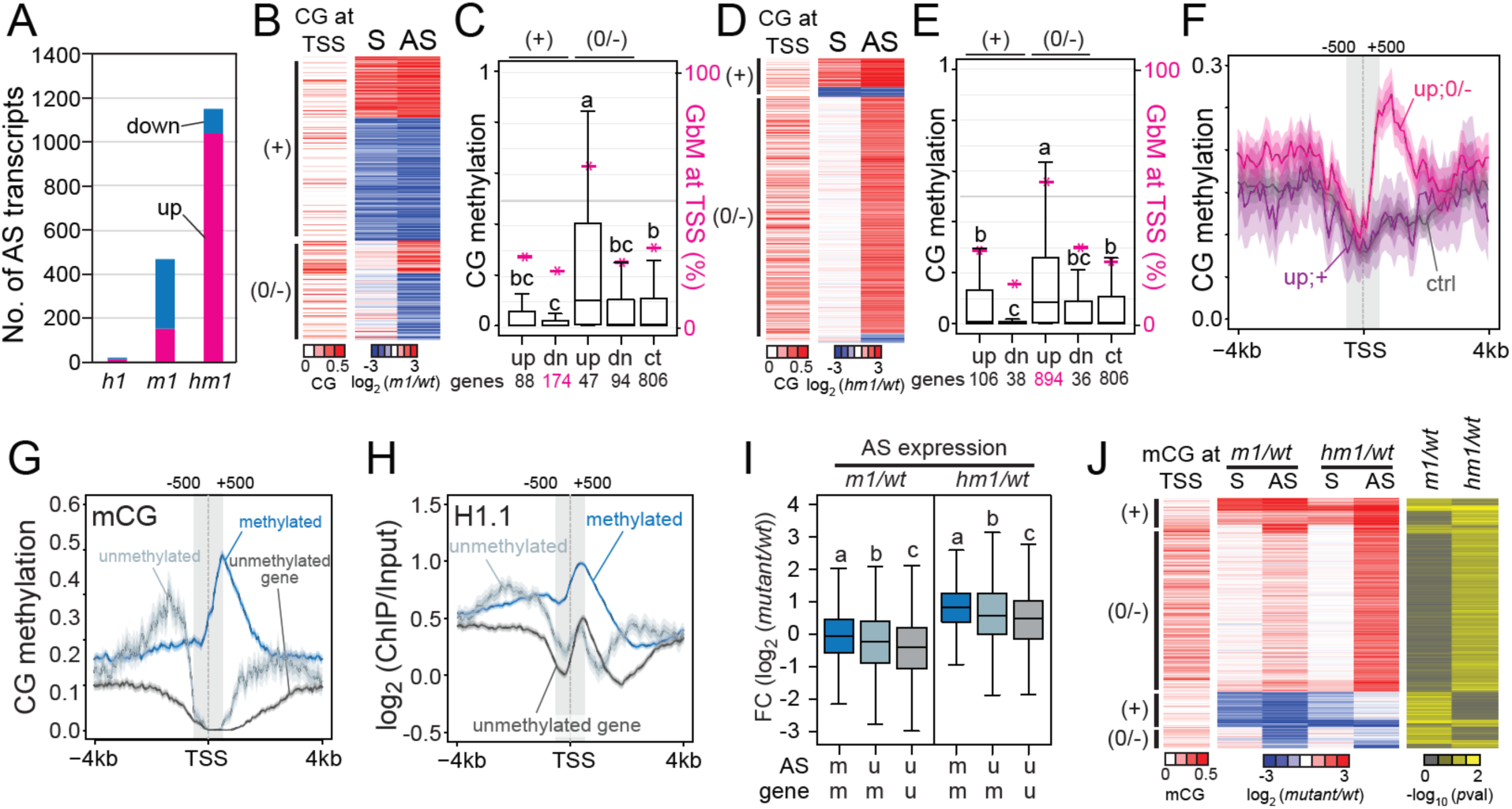
DNA methylation and H1 cooperatively suppress intragenic antisense transcripts. **(A)** Number of differentially expressed antisense (AS) transcripts in *h1, met1*, and *h1met1* plants. **(B and D)** Heat maps showing *wt* CG methylation at antisense TSS and expression of sense (S) and antisense transcripts in *met1* (B) or *h1met1* (D) plants. **(C and E)** Box plots of TSS-proximal *wt* CG methylation levels of upregulated (up) and downregulated (dn) antisense transcripts either positively correlated (+) or negatively/un-correlated (0/-) with sense expression in *met1* (C) or *h1met1* (E) plants. Data are also shown for unchanged (control; ct) antisense transcripts. Magenta asterisks indicate the percentage of antisense TSS that overlap gbM. “a”, “b” and “c” are significantly different; overlapping letters – i.e. “b” and “bc” – indicate that the difference is not significant (*p* < 0.01; ANOVA). **(F)** *Wt* CG methylation around antisense TSS of indicated groups from (E) with more than 100 transcripts. Shaded areas represent 95% confidence intervals. **(G-H)** CG methylation (G) and H1.1 distribution (H) around the TSS of antisense transcripts initiated from unmethylated genes, unmethylated regions within gbM genes, and methylated regions within gbM genes. **(I)** Box plots of antisense expression in indicated groups from (G) in *met1* and *h1met1* plants. “a”, “b” and “c” are significantly different (*p* < 0.01; ANOVA). **(J)** Heat maps of *wt* CG methylation at antisense TSS, expression of sense and antisense transcripts compared to *wt*, and corresponding antisense expression *p*-values in *met1* and *h1met1* plants. **(C, E and I)** Whiskers indicate 1.5X IQR.

We next separated antisense transcripts for which the change in expression is positively correlated with the sense transcript from those with an absent or negative correlation (some antisense transcripts were ambiguously correlated and therefore were removed from this analysis). A positive correlation suggests that antisense expression is altered due to increased or decreased overall transcriptional activity at the locus, whereas uncorrelated or negatively correlated antisense transcription must be regulated differently. In *met1* plants, sense and antisense expression is correlated in most cases (65%, 262 out of 403 transcripts in Figure 6B and Table S4), indicating that the antisense transcriptional changes we detected are mainly due to mis-regulation of sense transcripts. However, negatively or un-correlated, upregulated antisense transcripts are much more likely to initiate from methylated DNA than positively correlated, downregulated, or unchanged transcripts (Figures 6B and 6C). This result suggests that antisense expression is activated by methylation loss, but we identified just 47 (12%) such transcripts (Figures 6B and 6C). Thus, *met1* data provide only modest support for the hypothesis that gbM suppresses antisense expression.

The situation is substantially different in *h1met1* plants, in which antisense expression is most commonly upregulated and not positively correlated with sense expression (83%, 894 out of 1074 transcripts in Figure 6D and Table S4). As in *met1*, these transcripts are much more likely to initiate from DNA methylated in *wt* than other antisense transcripts (Figures 6D-F and S6A, S6B), indicating that gbM and H1 cooperatively silence antisense transcription.

To further analyze the link between gbM and antisense expression, we separated all the antisense transcripts we annotated (regardless of their behavior in mutant genotypes) into three groups based on their point of initiation: those from methylated DNA, from unmethylated DNA within methylated genes, or from unmethylated genes (Figure 6G). As expected, transcripts that initiate from methylated DNA have more H1 around the TSS (Figures 6H and S6C, S6D). In *met1*, levels of transcripts arising from methylated genes are significantly elevated compared to unmethylated genes (Figure 6I). Transcripts initiating from methylated sequences are also significantly elevated compared to those initiating from unmethylated regions of methylated genes (Figure 6I). In *h1met1*, levels of transcripts arising from methylated genes are also significantly elevated, and transcripts initiating from methylated DNA are especially upregulated (Figure 6I), specifically linking methylation loss with antisense transcript activation.

TEs that are highly active in the absence of H1 and DNA methylation tend to show weak upregulation when methylation alone is lost (Figures 5A and 5E). We therefore asked if antisense intragenic transcription is analogously affected. Indeed, antisense transcription tends to behave similarly in *met1* and *h1met1* (Figure 6J). Of particular interest, transcripts that are significantly upregulated only in *h1met1* are usually also activated to some extent in *met1* (Figures 6J and S6B). Thus, as with TEs, loss of H1 hyperactivates antisense transcripts that are released from silencing by elimination of intragenic methylation.

## Discussion

DNA methylation and histone H1 are abundant and widely conserved constituents of eukaryotic chromatin associated with transcriptional inactivity (Fyodorov et al., 2018; Torres et al., 2016; Zhang et al., 2018a), and their interactions have been of interest for many years. There is now extensive evidence that H1 modulates DNA methylation pathways. Loss of H1 causes genome-wide hypermethylation in ascomycete fungi (Barra et al., Mol Cell Biol 2000; Seymour, G3 2016) and reduces methylation at specific loci in mouse (Fan et al., 2005; Geeven et al., 2015; MacLean et al., 2011; Yang et al., 2013). In *Arabidopsis*, loss of H1 reduces methylation of euchromatic TEs and increases methylation of heterochromatic elements in all sequence contexts (Rutowicz et al., 2015; Zemach et al., 2013). H1 also appears to impede DNA demethylation in *Arabidopsis* heterochromatin (He et al., 2018). H1 hinders heterochromatic methylation by restricting the access of DNA methyltransferases (Lyons and Zilberman, 2017), and likely interferes with demethylation in an analogous way. The ability of H1 to globally influence nucleosome positions in plants (Figure 2) and animals (Baldi et al., 2018; Fan et al., 2003; Woodcock et al., 2006) suggests an additional regulatory mechanism, because nucleosomes are substantial obstacles to DNA methylation (Baubec et al., 2015; Huff and Zilberman, 2014; Lyons and Zilberman, 2017).

The effects of DNA methylation on H1 are far less established. A number of studies evaluated whether H1 has preferential affinity for methylated DNA *in vitro* but came to opposing conclusions (Campoy et al., 1995; Hashimshony et al., 2003; Levine et al., 1993; McArthur and Thomas, 1996; Nightingale and Wolffe, 1995). Our results show that H1 is enriched at methylated loci in *Arabidopsis*, but this enrichment does not depend on DNA methylation *in vivo* (Figure 1). Our data indicate that the regulatory relationship between H1 and DNA methylation is unidirectional: H1 modulates, but is not directly affected by, DNA methylation. However, methylation indirectly influences H1 by, for example, enforcing TE silencing (Figure 1F).

The complex relationship between H1 and DNA methylation ultimately impinges on chromatin accessibility and transcription. Our results indicate that H1 and DNA methylation jointly maintain heterochromatic TEs in an inaccessible and silent state (Figure 5). Methylation is the more powerful repressor, as few TEs are activated in *h1* mutants, and many of those that gain activity also lose methylation (Figure 4). However, the role of H1 becomes obvious when methylation is reduced, and loss of H1 can even overcome modest hypermethylation to activate TEs (Figure 5E). These results may explain why H1 depletion has a far more drastic effect on TE activity in *Drosophila* (Iwasaki et al., 2016; Lu et al., 2013), which lacks cytosine DNA methylation (Raddatz et al., 2013; Zemach et al., 2010), than in mouse (Fan et al., 2005).

The cooperative silencing of transcription by methylation and H1 extends to gene bodies (Figure 6). The function of gbM has been extensively debated for over a decade. One line of argument is that gbM reprogramming regulates development (Herb et al., 2012), for example by modulating alternative splicing (Lyko et al., 2010; Park et al., 2011). These ideas are supported by developmental disruptions caused by erasure of methylation and/or downregulation of DNA methyltransferases in species in which methylation is exclusive or near-exclusive to gene bodies (Bewick et al., 2018; Biergans et al., 2017; Kucharski et al., 2008). However, in every species examined so far, gbM is concentrated in constitutively expressed housekeeping genes (Coleman-Derr and Zilberman, 2012; Dixon et al., 2016; Sarda et al., 2012; Zilberman, 2017). These genes do not vary in expression during development and thus are not good candidates for differential alternative splicing or other developmental regulation. There is also little evidence for substantial shifts in gbM patterns during plant or animal development (Bartels et al., 2018; Libbrecht et al., 2016; Suzuki et al., 2013; Zemach et al., 2018). Another argument is that gbM is a functionless consequence of TE methylation (Bewick and Schmitz, 2017; Bewick et al., 2016; Teixeira and Colot, 2009), motivated by the inability to detect transcriptional or RNA processing changes caused by genetic removal of gbM. This argument is undermined by the well-established observation that methylation is mutagenic in plants and animals (Alexandrov et al., 2013; Takuno and Gaut, 2013). The maintenance of DNA methylation within the exons of some of the most conserved and essential genes, especially in species that do not methylate other sequences, implies a function that is important enough to outweigh the costs of increased mutation rates (Hunt et al., 2013; Takuno and Gaut, 2013).

The last line of argument is that developmentally invariant methylation within housekeeping genes should have one or more housekeeping functions (Zilberman, 2017). This is supported by observations of small overall drops in gene expression associated with evolutionary loss of gbM in plants (Muyle and Gaut, 2018; Takuno et al., 2017). One housekeeping function, the inhibition of intragenic aberrant transcripts, was proposed when gbM was first discovered (Tran et al., 2005). Mechanisms to silence such transcripts have been described in multiple species, indicating that this is an important feature of transcriptional homeostasis (Carrozza et al., 2005; Kaplan et al., 2003; Neri et al., 2017; Whitehouse et al., 2007). Here we provide the long-sought experimental evidence for this gbM function. Our data show that intragenic antisense transcription is upregulated in *h1met1* plants (Figure 6A). Most upregulated transcripts are anticorrelated with sense expression (Figure 6D) and about half of these (55%) initiate from methylated DNA – far more than is expected by random chance (Figure 6E). Conversely, antisense transcripts that initiate from methylated DNA are preferentially upregulated in *h1met1* (Figure 6I). Antisense transcripts activated in *h1met1* also tend to be upregulated, albeit more weakly, in *met1* (Figure 6J). Taken together with the observed silencing of TEs by methylation and H1, the suppression of gene transcription by H1, and the ability of H1 to associate with chromatin independently of DNA methylation, our results strongly support the hypothesis that H1 and DNA methylation cooperatively repress aberrant intragenic transcripts in *Arabidopsis*. The strong similarities between gbM patterns of plants and animals (Feng et al., 2010; Zemach et al., 2010), and the abundance of H1 in animal genes (Cao et al., 2013; Millán-Ariño et al., 2014), suggest that this function is general, and potentially universal among eukaryotes with intragenic methylation.

## Acknowledgements

We thank X. Feng, E. Hollwey and S. de-Bruijin for helpful comments on the manuscript. We also thank S.L. McDevitt and M. Chung for sequencing and C. Wistrom for greenhouse assistance.

## Author contributions

J.C. designed and performed experiments, analyzed data and wrote the manuscript. D.B.L. performed cytology, image analysis, NRL calculation and MNase-seq analysis. M.Y.K. performed H1 ChAP-seq experiments. J.D.M. defined the boundary of gbM regions. D.Z. conceived and supervised the project and wrote the manuscript.

