## Supplemental Information for "DNA methylation and histone H1 cooperatively repress transposable elements and aberrant intragenic transcripts"

### Methods

#### Biological materials

The *met1-6* (*met1*) (Xiao et al., 2003) and *h1.1 h1.2* (*h1*) (Zemach et al., 2013) mutant lines were described previously. To establish *h1.1 h1.2 met1-6* triple mutants (*h1met1* mutants), *h1.1 h1.2* plants were crossed with *met1-6 +/-* heterozygous mutants. *h1.1 h1.2 met1-6 +/-* plants were isolated at F2 generation. F3 segregants were genotyped to identify homozygous triple mutants. F4 homozygous *h1met1* plants were used for this study. For ChIP-seq, RNA-seq, and bisulfite-seq samples, plants were germinated and grown for 2 weeks in half strength Gamborg's B-5 liquid media (Caisson Labs, cat. No. GBP07) under continuous light with shaking at 125 rpm. For ChAP-seq samples*, pH1.1-H1.1* and *pH1.2-H1.2* constructs were fused with biotin ligase recognition peptide (BLRP) at the N-terminus, then transformed into Col-0 plants (Zilberman et al., 2008). The transgenic plants were grown in liquid culture for 4 weeks.

#### Bisulfite sequencing (BS-seq) library preparation

Genomic DNA (gDNA) was isolated from 2-week-old seedlings using DNeasy plant mini kit (Qiagen, cat. No. 69104) according to the manufacturer's manual. About 500 ng of purified gDNA were fragmented to roughly 100-1000 bp then purified using 1.2X volume of SPRI beads (Beckman Coulter, cat. No. A63881). Fragmented gDNA was end-repaired and ligated to HPLC/PAGE purified, cytosine-methylated Illumina adapters (Elim Biopharmaceuticals, Inc.). DNA was subjected to two rounds of bisulfite conversion according to manufacturer's protocol (Qiagen, cat. No. 59104). To remove self-ligated adapters, DNA was purified twice with 0.8X volume of SPRI beads. NEB next indexing primers were used to amplify bisulfite-converted libraries (New England BIolabs Inc. cat. No. E7335S).

**Chromatin immunoprecipitation (ChIP) sequencing library preparation**

For H3K9me2 ChIP-seq libraries, 2 g of 2-weeks-old seedlings were cross-linked with 32 ml of crosslinking buffer (0.4 M sucrose, 10 mM tris-HCl pH 8.0, 1 mM EDTA and 1% formaldehyde (Thermo Scientific, cat. No. 28906)) for 10 min under vacuum. Cross-linking was quenched by applying vacuum for 5 min with 2 ml 2 M glycine. Cross-linked seedlings were washed with distilled water, pat dried with paper towels, frozen and ground in liquid nitrogen with pestle and mortar. Ground tissues were re-suspended in 20 ml of nuclei isolation buffer at 4 °C (NIB; 0.25 M sucrose, 15 mM PIPES pH 6.8, 5 mM $MgCl_2$, 60 mM KCl, 15 mM NaCl, 1 mM $CaCl_2$, 0.9% Triton X-100, 1 mM PMSF, 1X complete EDTA-free protease inhibitor cocktail (Roche, cat. No. 11873580001)). Homogenized slurry was filtered through Miracloth (Millipore, cat. No. 475855) and centrifuged at 6,000 rpm for 20 min at 4 °C to pull-down nuclei. After discarding supernatant, cell pellets were re-suspended in 1 ml 4 °C nuclei lysis buffer (NLB; 50 mM HEPES pH 7.5, 150 mM NaCl, 1 mM EDTA, 0.5 % SDS, 1 % Triton X-100, 1X complete EDTA-free protease inhibitor). 500 µl nuclei solution was sonicated and centrifuged for 10 min at 13,000 rpm in table-top centrifuge (4 °C). Sonicated chromatin in supernatant was transferred and diluted 5 times with dilution buffer (NLB without SDS and Triton X-100). 200 µl of diluted chromatin was incubated with 7 µl of antibodies against H3K9me2 (Abcam, cat. No. AB1220) overnight with rotation at 4 °C. 45 µl of pre-cleared protein G dynabeads (Invitrogen, 10003D) were added and incubated for 4 hours with rotation at 4 °C. 20 µl of diluted chromatin solution was kept for input control. Chromatin bound to the antibodies was washed 4 times with low salt wash buffer (150 mM NaCl, 20 mM Tris-HCl pH 8.0, 0.2% SDS, 0.5% Triton X-100, 2 mM EDTA) and one time with TE buffer (1 mM EDTA and 10 mM Tris-HCl pH 8.0) at 4 °C. Washed chromatin was re-suspended in 200 μl elution buffer (1% SDS), and incubated at 65 °C for 5 hrs with 8 μl of 5M NaCl (vortex for 1 hr). To digest proteins bound to gDNA, samples were incubated 1 hr at 55 °C with 4 μl protease K (Thermo, cat. No. EO0492), 4 μl of 0.5 M EDTA, 8 μl of 1 M tris-HCl pH 8. After reverse-crosslinking, 220 μl Phenol:Chloroform:Isoamyl alcohol 25:24:1, pH 8.0 (P:C:I; Sigma, cat. No. P3803), 500 μl ethanol, 20 μl 3M sodium acetate (pH 5.2), and 2 μl glycogen were added and incubated overnight at -80 °C to extract DNA. Precipitated DNA was washed with 70% ethanol and diluted in Milli-Q water. Isolated gDNA was subject to Illumina library construction as described in BS-seq library preparation without two rounds of bisulfite conversion.

**Chromatin Affinity purification (ChAP) sequencing library preparation**

ChAP experiments were performed according to previous publication with minor modifications (Zilberman et al., 2008). 4-week-old roots were vacuum infiltrated in 1% formaldehyde solution for 15 minutes to cross-link the chromatin. Roots were ground in liquid nitrogen and nuclei were isolated as described (Zilberman et al., 2008). The nuclear suspension was fragmented by sonication to 0.5-2 kb, then 90% of biotinylated protein-containing chromatin was affinity purified using streptavidin beads. 10% of the sheared chromatin suspension was used as input control. Reverse cross-linking was performed with 5M NaCl, and the DNA was purified as described (Zilberman et al., 2008). Isolated gDNA was subject to Illumina library construction as described in ChIP-seq library preparation.

**DNase I sequencing library preparation**

*Arabidopsis* nuclei were isolated and re-suspended in NIB as described in ChIP-seq library preparation. After centrifugation at 6,000 rpm for 20 min at 4 °C, nuclei pellets were re-suspended in 400 μl DNase I buffer (10 mM Tris-HCl pH 8.0, 2.5 mM MgCl2, 0.5 mM CaCl2). 100 μl of nuclei was incubated with 0.1-2 U / ml DNase I (New England Biolabs, cat. No. M03030S) for 3 min at 37 °C. 50 mM EDTA, 0.5% SDS (final concentration), 2 μl protease K were added and incubated at 55 °C for 1 hr to stop digestion. To completely denature chromatin, more SDS was added to a final 1% concentration then incubated at 65 °C for 15 min. gDNA was isolated with P:C:I and DNA fragmentation was visualized by DNA electrophoresis. Samples with similar degree of DNA digestion were selected to construct libraries. 0.5 X volume of SPRI beads were added to remove undigested gDNA. Supernatant was collected and purified with 1.25 X volume of SPRI beads. Isolated DNA was subject to Illumina library construction as described in ChIP-seq library preparation.

**RNA sequencing library preparation**

Total RNA was extracted from 2-week-old seedlings using Trizol (Invitrogen, cat. No. 15596026) according to manufacturer's manual. 1 μg of RNA was subjected to DNA-*free* DNA removal kit (Thermo, AM1907) to degrade DNA from samples. To remove rRNA, RNA was treated with Ribo-zero plant kit (Illumina, MRZPL1224). 50 ng of rRNA-depleted RNA was used to construct RNA sequencing library (Epicentre, cat. No. SSV21124) following the manufacturer's manual except for the RNA fragmentation step (90 °C, 10 min).

**Sequencing libraries**

All of sequencing in this research was performed at the Vincent J. Coates Genomic Sequencing Laboratory at the University of California, Berkeley with HiSeq 2500 or HiSeq 4000.

### Nuclear cytology

For DAPI staining of nuclei, whole 2-week old *Arabidopsis* seedlings grown on B5 agar plates with continuous light were fixed in 4% PFA in PBS at room temperature under a vacuum for 30 minutes. Three 30-minute PBS washes were then carried out and seedlings were subsequently incubated with 0.1% DAPI in PBS and 0.05% Tween-20 rocking in a 12-well plate at room temperature for 1 hour. Following 2 PBS washes, seedlings were squash-mounted on slides in Vectashield and imaged.

### Image analysis

Laser scanning confocal microscope z-stacks were acquired using a Zeiss LSM710. To generate 2D representations of these images, stacks were converted into max Z-projections. These projections were then fed into Fiji function "3D surface plot" (Schindelin et al., 2012) to generate 3D DAPI intensity renderings. Imaris software (Bitplane) was used to visualize heterochromatic puncta with the "surfaces" option (minimum feature size =0.07 μm for identifying chromocenters; set to 0.5 μm for whole nucleus rendering) as well as to calculate the approximate volume of these puncta.

### Sequence alignments

For RNA-seq libraries, reads were mapped to TAIR10 genome using HISAT2 (Kim et al., 2015; Pertea et al., 2016) with following options: --max-intronlen 10000 -k 2 --dta. Only uniquely mapped reads were kept for further analysis. For BS-seq libraries, reads were mapped with bs-sequel pipeline (https://zilbermanlab.net/tools/). For other sequencing libraries, reads were mapped with Bowtie (Langmead et al., 2009) allowing up to 2 mismatches. Samples were normalized by reads per kilobase per million mapped reads (RPKM) when input samples were available. Otherwise, samples were normalized by bins per million mapped reads (BPM). Treated samples were divided by input samples (when input samples were available) to calculate enrichment, and transformed into log 2 values using bamcompare pipeline in deepTools2 (bamcompare --scaleFactorsMethod None –normalizeUsing RPKM, or bamcoverage –normalizeUsing BPM) (Ramírez et al., 2016). We used 'window_alignment' and 'gff_arithmetics' Perl scripts (https://zilbermanlab.net/tools/) and awk to calculate the log 2 ratio of treatment and input for DNaseI-seq libraries.

**Re-annotation of transposable element (TE) genes**

Most TE genes are silenced in *wt* plants. Because the Araport11 annotation was based on the RNA expression in 113 RNA-seq data generated in Col-0 *wt* plants (Cheng et al., 2017), TE annotations are more likely to rely on predictions compared to gene annotations. This encouraged us to re-annotate expressed TE genes using our dataset.

Aligned reads were separated by their directionality (Watson and Crick strands). Biological replicates were merged to increase sensitivity and accuracy of annotation. StringTie (Pertea et al., 2015, 2016) was accommodated to re-annotate expressed TEs with guided reference for Araport11 genes with following options: -g 300 --fr -t -c 0.5 -m 110. Annotations from each dataset were merged into a single annotation using 'stringtie –merge' tool (options: -F 0 -T 0.5 -f 0.1).

To identify TE genes, 1364 methylated (mCG and mCHG > 0.1, mCHH > 0.02), heterochromatic (H3K9me2 > 0 (log2(ChIP/Input)), and expressed (transcripts per kilobase per million reads (TPM) > 5) elements were isolated (Table S1). 72% of these (987/1364) overlapped with Araport11 TE annotation (gffcompare in StringTie package; any class code except for 'p' and 'u', which indicates 3' run off transcripts and no overlap, respectively) (Pertea et al., 2015, 2016). Araport11 TE gene annotations were replaced with our StringTie TE gene annotation if there was any overlap, except for 72 StringTie TE gene annotations that exactly matched to Araport11 annotation.

#### *De novo* annotation of antisense transcripts

Using our StringTie annotations, any transcripts that overlaps with Araport11 genes (mCHG < 0.01, mCHH < 0.01) in antisense direction ('x' and 's' in gffcompare) were isolated (Pertea et al., 2015, 2016).

#### Description of *Arabidopsis* genome features

'Genes' indicate representative gene annotations from Araport11 (Cheng et al., 2017), excluding ones with more than 1 % CHG or CHH methylation within gene bodies in *wt* plants. 'Transposable elements' include TE, TE genes and pseudogenes longer than 250 bp in Araport11, including 1292 re-annotated TE genes in this study (Table S1). 'Heterochromatic transposons (hTEs)' refers to methylated (mCG and mCHG > 0.1, mCHH > 0.02) heterochromatic (H3K9me2 > 0 ($log_2$(ChIP/Input)) TEs. 'Euchromatic transposons' indicates methylated TEs with low H3K9me2 (H3K9me2 < 0 ($log_2$ (ChIP/Input)). We identified *wt Arabidopsis* 'gene-body methylated regions' using a modified version of the R package MethylSeekR (Burger et al., 2013) to analyze triplicate S3 lines from a previous epimutation accumulation experiment (Schmitz et al., 2011). The modification we made to MethylSeekR was to use mean segment methylation rather than segment length as the threshold to differentiate between unmethylated and methylated segments, as we found this to work better for *Arabidopsis*, in contrast to the mammalian models the package was originally designed for. We segmented combined CHG and CHH methylomes from the three lines to identify segments of 3 or more consecutive unmethylated non-CG cytosines. We then extracted corresponding segments from the combined CG methylome of the three lines and segmented these data to differentiate between fully unmethylated and mCG segments. In each case, we chose segmentation parameters by iterating through values of 'm' (the segment methylation threshold) and n (the minimum segment length). For each n, we plotted a ROC curve by varying m and comparing the overlaps of resulting putative methylated and unmethylated segments with sets of annotated transposons and annotated unmethylated genes. We chose values of n to maximize AUC using a trapezium approximation (n=500 for non-CG methylation, n=9 for mCG), and values of m to provide best separation between methylated and unmethylated segments by analysis of density distributions of segment mean methylation (m=0.15 for non-CG methylation, m=0.0931 for mCG). We observed that these segmentation parameters led to most mCG segments falling within gene bodies and comprising a single or small number of segments per gene. Having thus identified longer mCG segments (>=9 consecutive CG sites), we added shorter mCG segments within otherwise unmethylated regions by repeating the segmentation of these regions and reducing n (n=3; n=1). We then trimmed segments identified as mCG to exclude regions which fall outside the set of gene loci in the Araport11 genome annotation (Cheng et al., 2017).

**Identification of differentially methylated transcription start sites in TE genes**

Numbers of methylated cytosines and unmethylated cytosines at first 500 bp of expressed TE genes were counted by window_by_annotation Perl script (https://zilbermanlab.net/tools/). TE genes with $p$-value < 0.05 and methylation change greater than 10% (mCHG) or 2% (mCHH) were considered differentially methylated (Fisher's exact test). TE genes with *p*-value greater than 0.05 or methylation difference below 2% were considered not differentially methylated.

**Classification of MET1-dependent and independent hTEs**

MET1-dependent hTEs were defined as hTEs that lost H3K9me2 in *met1* knockout plants. To identify these TEs, we extracted average H3K9me2 level of 50 bp windows that overlap hTEs. hTEs containing windows that significantly lost H3K9me2 in *met1* plants compared to *wt* ($p < 0.05$, metilene (Jühling et al., 2016)) were isolated. To filter out TEs with local, but not global, reduction in H3K9me2, the average level of H3K9me2 is also considered ($H3K9me2_{met1} < H3K9me2_{wt}$). MET1-indpendent hTEs were defined as: Do not overlap with regions that significantly lost H3K9me2 in *met1* (*vs. wt*) and average $H3K9me2_{met1} > 0$.

**Identification of differentially expressed transcripts**

RNA-seq reads were pseudo-aligned to Araport11 genes and the abundance of transcripts was quantified using Kallisto (Bray et al., 2016). Differentially expressed genes were identified by Sleuth ($p < 0.05$) (Pimentel et al., 2017). Due to duplication of TE genes, pseudo-alignment based quantification can be less accurate to measure the expression of TE genes. Also, annotation of expressed TEs and antisense transcripts was based on the mapped RNA-seq reads. Therefore, we accommodated an alignment-based quantification tool (DeSeq2, (Love et al., 2014)) to identify differentially expressed TE genes and antisense transcripts (FDR < 0.1).

**Predicting H1 distribution in *wt* and *met1* plants**

Average DNA methylation and gene expression levels for 1 kb windows were transformed into log2. Windows without any DNA methylation were discarded then methylation level was transformed into log2. Log-transformed average GC content, DNA methylation and levels of other chromatin features, including histone H1, were centered to 0 and scaled between -0.5 to 0.5. To minimize bias from outliers, windows between the $5^{th}$ and the $95^{th}$ percentile were used for scaling data.

To visualize the relationship among genomic features, principal component analysis was applied to arrays of features using Gene Cluster 3.0 (de Hoon et al., 2004). First (PC1) and second (PC2) principal components were plotted in Figure 1H.

GC content, H3K9me2 and H3K4me3 were incorporated into a H1 prediction model to explain H1 distribution in *wt* and *met1* plants. Linear regression of H1 using averaged H1 ((H1.1+H1.2)/2), GC content, H3K9me2, and H3K9me3 distribution in *wt* plants was performed using the 'lm' function in R. The resulting model was:

$$H1_{wt} = 0.82 * (\text{GC content}) + 0.11 * (H3K9me2_{wt}) - 0.38 * (H3K4me3_{wt}) + 0.03$$

The H1 distribution in *met1* plants was predicted using the coefficients from the $H1_{wt}$ model and the distributions of H3K9me2 and H3K4me3 in *met1*.

**Clustering and heatmaps**

For clustering RNA expression data, TPM was calculated first, then the data for each gene was centered and normalized using Gene Cluster 3.0 (de Hoon et al., 2004). Average level of chromatin features (histone modifications, histone variants, DNA accessibility) and DNA methylation data was used without modifications. hTE clusters in Figure 5A and 5E were created as follows:

Differentially expressed hTEs in *met1* or *h1met1* were isolated (*vs. wt*, FDR < 0.1). hTEs with differentially methylated TSS in *h1met1 vs. met1* were separated from ones without methylation change (in CHG or CHH context). Each hTE group was clustered by *k*-means clustering with 2 to 5 clusters, then the number of clusters that consistently appeared as major clusters was determined (2 for hTEs without methylation change, 3 for hTEs with methylation change) (de Hoon et al., 2004). TEs that do not follow the expression pattern of their assigned cluster were removed. Clusters were visualized by TreeView (de Hoon et al., 2004). $Log_2$ expression fold change calculated by DeSeq2 was used to cluster antisense transcripts (Figures 6B, 6D and 6J). Clustering was performed as above. Positive correlation indicates: Both sense and antisense transcripts are upregulated or downregulated in mutants (*vs. wt*; $log_2FC > 0.5$ or $< -0.5$, and *p*-value < 0.05 for sense transcripts (Sleuth) (Pimentel et al., 2017) and FDR < 0.1 for antisense transcripts (DeSeq2) (Love et al., 2014). No correlation or negative correlation indicates: When antisense transcripts are upregulated in mutants (*vs. wt*; $log_2FC > 0.5$ and FDR < 0.1), sense transcripts show no expression change or downregulated ($log_2FC < 0.2$), and *vice versa*.

**MNase-seq plotting**

Normalized smoothed bigwig files of WT MNase-seq were used in plotting nucleosomes anchored at well-positioned *wt* nucleosomes (Lyons and Zilberman, 2017) (GEO accession GSE96994).

**Nucleosome repeat length calculation**

Sequences of MNase-digested *Arabidopsis* chromatin (Lyons and Zilberman, 2017) were used to compute dyad locations with the nucleosome mapping software NuMap (Valouev et al., 2011). Output files were imported to R for NRL calculation and visualization (detailed scripts at https://github.com/dblyons/MNase-seq). Fragments 120 to 180 bp were filtered from MNase-seq libraries mapped to TAIR10 genome build. Nucleosome dyads were called, which were subsequently used to calculate a peak-to-peak phasogram. The peaks of this phasogram were fed into the built-in R function "lm" (linear model) to estimate the NRL (Valouev et al., 2011). NRL subsetting by H3K9me2, H1 ChIP, and gene expression level was accomplished by intersecting against appropriate genomic coordinates using IntersectBed (Quinlan and Hall, 2010).

**Data visualization**

Screenshot of *Arabidopsis* chromosome features were taken by IGV (Robinson et al., 2011; Thorvaldsdottir et al., 2013). Averaged H1 and DNA methylation distributions between TSS and TTS were generated with Seqplots (Stempor and Ahringer, 2016). Average enrichment score matrices of genomic features of interest were generated by Perl scripts (https://zilbermanlab.net/tools/), then imported to R (Davey et al., 1997) for downstream analysis including boxplots and scatter plots.

**Use of previously published data**

Published data for histone variants and histone modifications were used to predict H1 distribution in *Arabidopsis*. Histone H2A.W data (Yelagandula et al., 2014), *wt* MNase-seq data (Lyons and Zilberman, 2017), and histone modification data (H3K9ac, H3k27ac (Chen et al., 2017), H3K36me2, H3K36me3 (Luo et al., 2013)) were obtained through GEO (GSE50942, GSE96994, GSE79524, and GSE28398). H3K4me3 data (Choi et al., 2018) were downloaded from ArrayExpress (E-MTAB-5048).

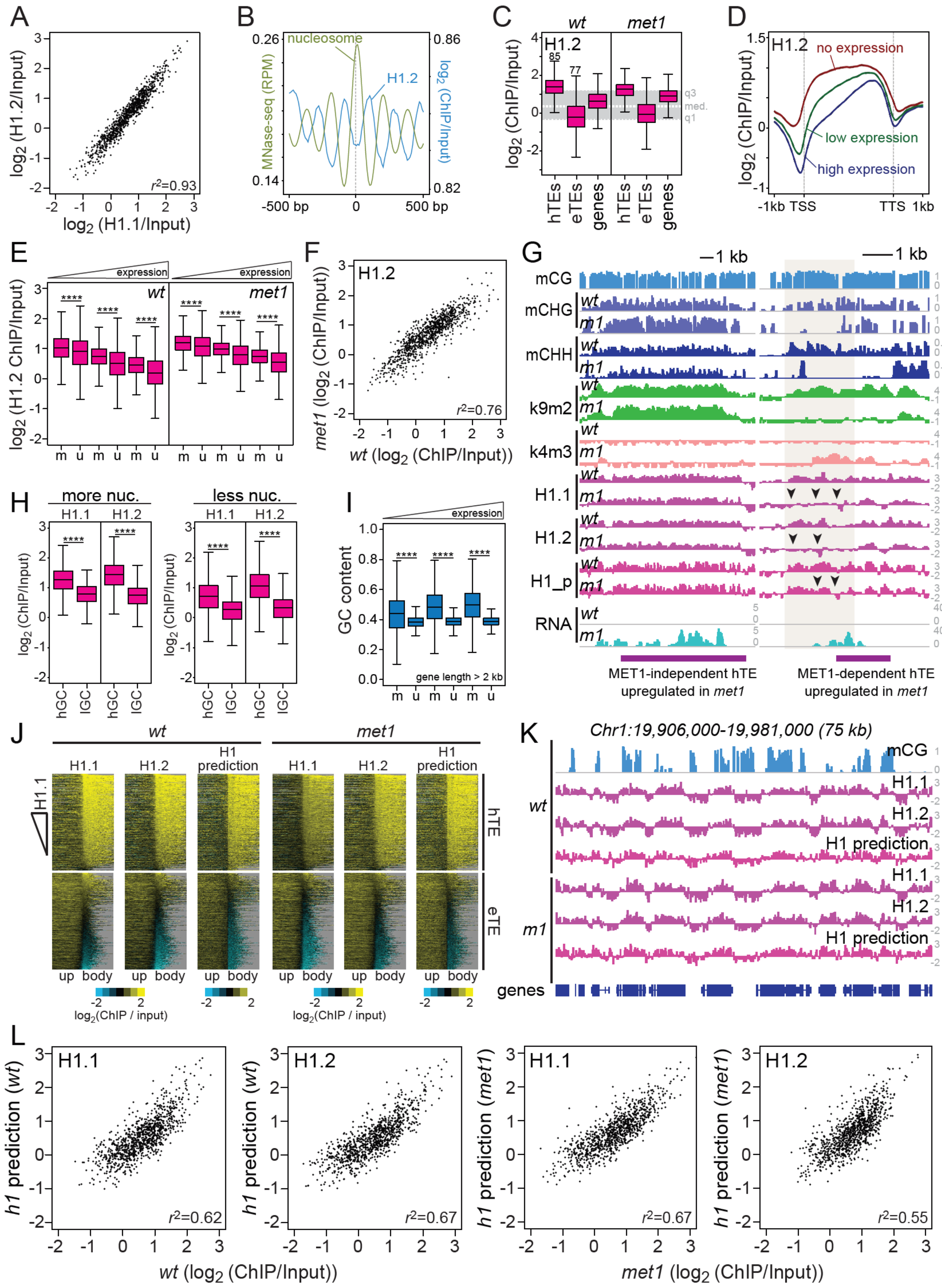
A
B
C
D
E
F
G
H
I
J
K
L
nucleosome
H1.2
MNase-seq (RPM)
no expression
low expression
high expression
expression
wt
met1
more nuc.
less nuc.
gene length > 2 kb
MET1-independent hTE upregulated in met1
MET1-dependent hTE upregulated in met1
Chr1:19,906,000-19,981,000 (75 kb)
H1 prediction
log2(ChIP / input)
h1 prediction (wt)
h1 prediction (met1)
wt (log2 (ChIP/Input))
met1 (log2 (ChIP/Input))

**Figure S1. Histone H1 distribution can be explained by GC content and features of repressive and active chromatin, Related to Figure 1. (A)** Average H1.1 and H1.2 levels of 1 kb windows in *wt*. 1000 windows were randomly selected for plotting. **(B)** Average H1.2 distribution around well-positioned nucleosomes. **(C)** Box plots of H1.2 levels in hTEs, eTEs, and genes. The shaded area marks the middle 50% of genomic H1.2 levels. Numbers (85, 77) indicate average CG methylation at hTEs and eTEs, respectively. **(D)** H1.2 distribution around genes with different expression levels. **(E)** Boxplots of H1.2 abundance in methylated (m) and unmethylated (u) genes in *wt* and *met1* plants. **(F)** Average H1.2 levels of 1 kb windows in *wt* and *met1*. 1000 windows were randomly selected for plotting. **(G)** Examples of actual H1 and predicted H1 distribution at a MET1-independent hTE (*AT2G12210*) and a MET1-dependent hTE (*MSTRG.47084*, Chr5:9206449-9207752) in *wt* and *met1*. Black arrows indicate regions of H1 loss in *met1* plants. **(H)** Box plots of H1 levels in nucleosome-rich (more nuc.) and nucleosome-poor (less nuc.) genes with high (hGC) or low (lGC) GC content ((G+C)/(A+G+C+T)). **(I)** Box plots of GC content in methylated (m) and unmethylated (u) genes. **(J)** Linear model prediction of H1 distribution around the 5’ end of TEs in comparison to actual H1 distributions. **(K)** An example of actual H1 and predicted H1 distribution at an array of gbM genes in *wt* and *met1* plants. **(L)** Average actual H1 and predicted H1 levels of 1 kb windows in *wt* and *met1*. 1000 windows were randomly selected for plotting. **** is $p < 0.0001$, Student’s *t*-test. “*r*” is Pearson’s correlation coefficient. **(C, E, H and I)** Whiskers indicate 1.5X IQR.

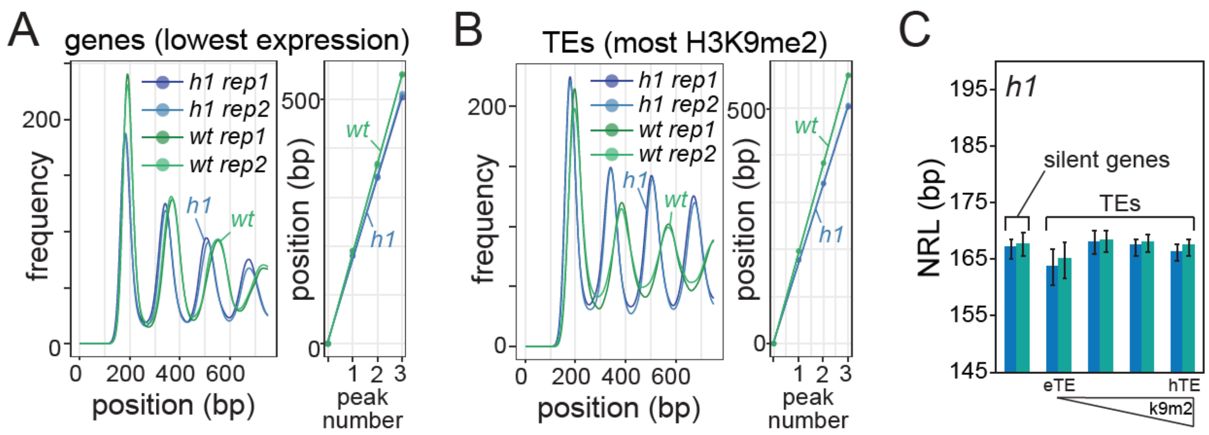


**Figure S2. Loss of H1 alters nucleosome organization in genes and TEs, Related to Figure 2.** **(A)** Phasograms and linear regressions for the positions of nucleosome peaks at the genes with lowest expression (shaded area in Figure 2A) in *wt* and *h1* plants. **(B)** Phasograms and linear regressions for the positions of nucleosome peaks at H3K9me2-rich TEs (shaded area in Figure 2D) in *wt* and *h1* plants. (C) NRL calculations for two biological replicates of *h1* plants at silent genes and TEs grouped by their H3K9me2 level in *wt*. Error bars indicate standard error.

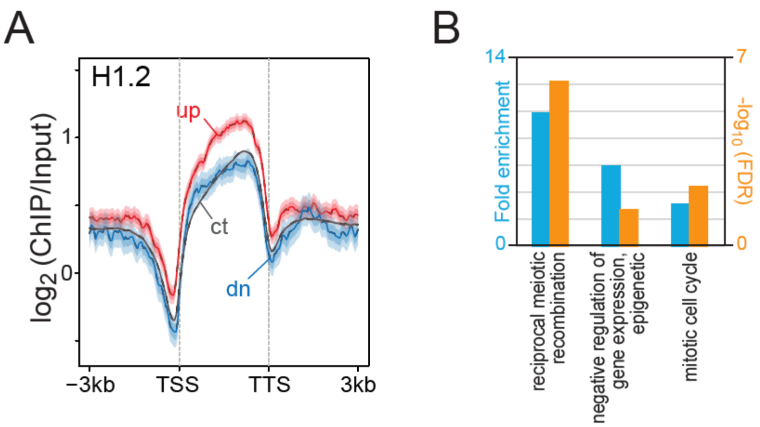


**Figure S3. H1 distribution and enriched gene ontology terms for genes mis-regulated in *h1* plants, Related to Figure 3. (A)** Distribution of H1.2 around control, up- and down-regulated genes. Shaded areas represent 95% confidence intervals. **(B)** Fold enrichment and corresponding false discovery rate (FDR) for the three enriched gene ontologies among up-regulated genes in *h1* plants (vs. *wt*).

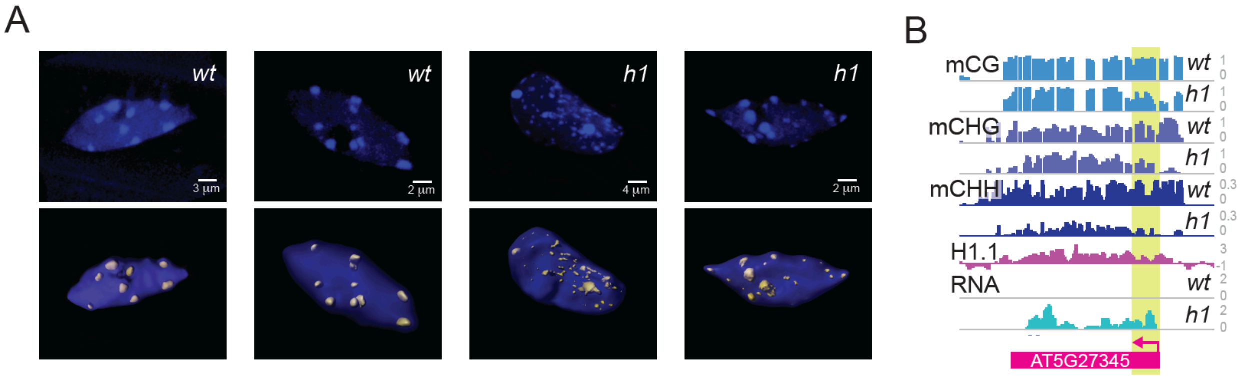


**Figure S4. Loss of H1 disperses heterochromatin and activates TEs, Related to Figure 4. (A)** Nuclei of *wt* and *h1* plants stained with DAPI. DAPI puncta volumes are highlighted in yellow. **(B)** An example of upregulated TE in *h1* plants. Note DNA methylation loss near TSS (highlighted in yellow).

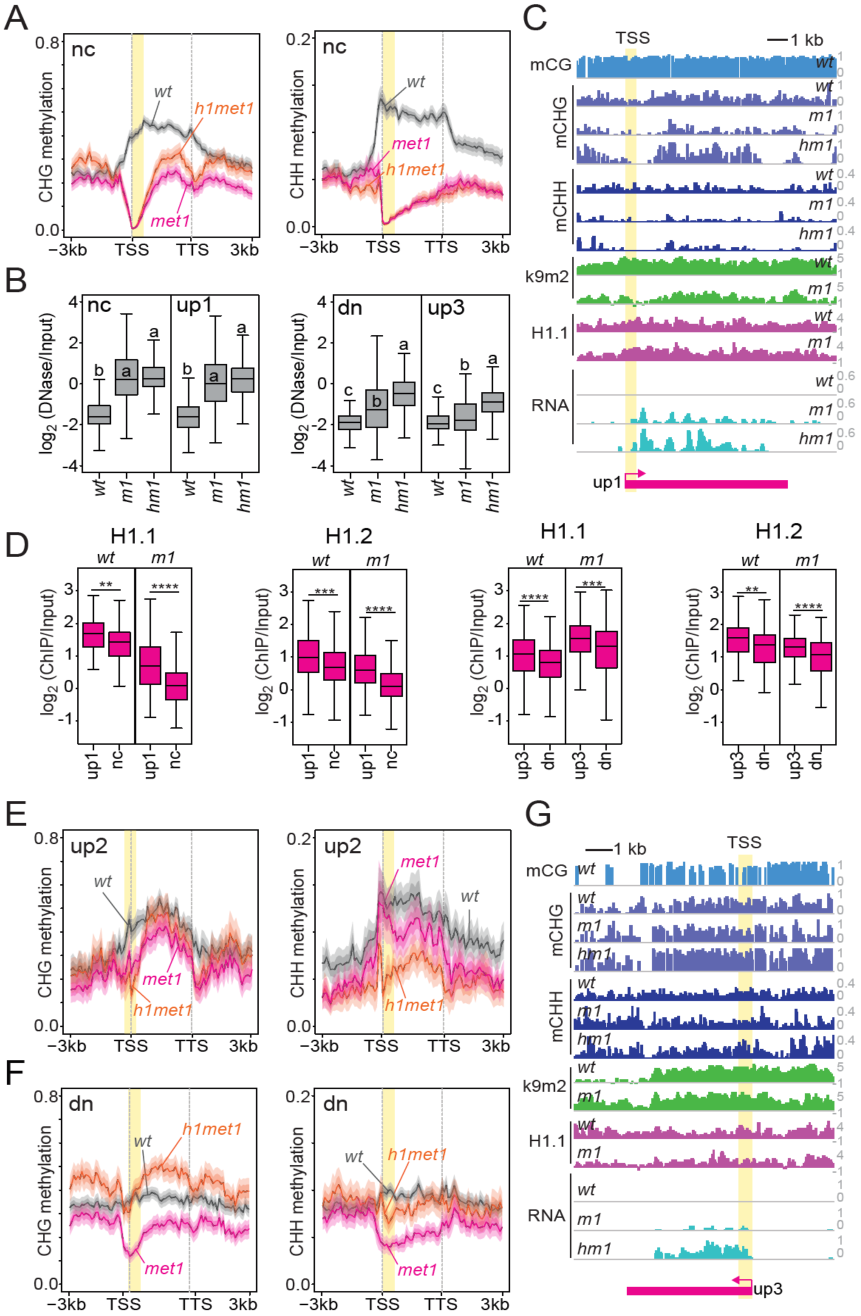


**Figure S5. Non-CG methylation, DNA accessibility and TE expression in *wt, met1* and *h1met1* plants, Related to Figure 5. (A)** Average CHG and CHH methylation around TEs in nc cluster from Figure 5A. **(B)** Box plots of DNA accessibility of TEs in clusters from Figure 5A and 5E in *wt*, *met1* and *h1met1* plants. “a”, “b” and “c” are significantly different ($p < 0.01$; ANOVA). **(C and G)** Examples of DNA methylation, H1.1 distribution and transcription at TEs in up1 (*AT3G33193*; C) and up3 (*AT1G40075*; G) clusters. **(D)** Box plots of *wt* and *met1* (*m1*) TE H1 levels in clusters from Figure 5A and 5E. ** is $p < 0.01$, *** is $p < 0.001$, and **** is $p < 0.0001$, Student’s *t*-test. **(E-F)** Average CHG and CHH methylation around TEs in up2 (E) and dn (F) clusters from Figure 5E. **(B, D)** Whiskers indicate 1.5X IQR.

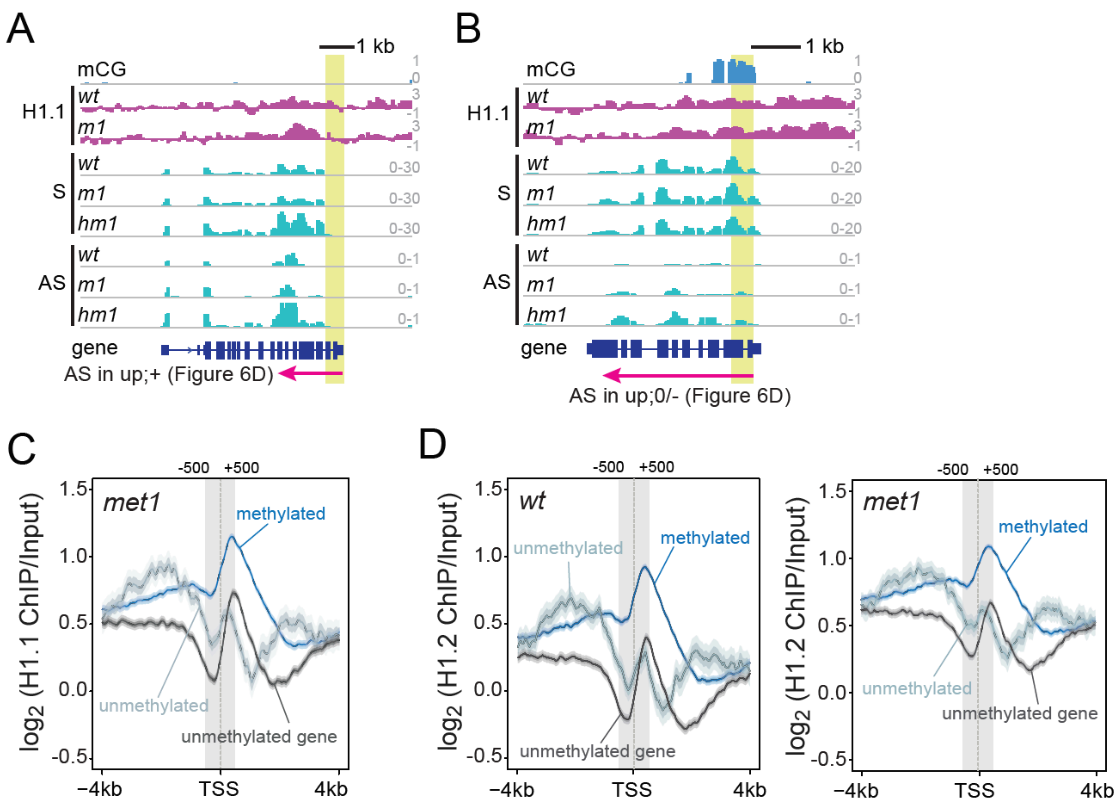


**Figure S6. CG methylation and histone H1 at antisense-transcribed genes, Related to Figure 6. (A-B)** Examples of CG methylation, H1.1 distribution, and sense and antisense expression at antisense transcripts either positively (A, up;+, gene *AT2G47240*) or negatively/un-correlated (B, up;0/-, gene *AT2G47240*) with sense expression. **(C-D)** H1 distribution around the TSS of antisense transcripts in *wt* and *met1* plants initiated from unmethylated genes, unmethylated regions within gbM genes, and methylated regions within gbM genes.
